# Skeletal muscle stem cell self-renewal and differentiation kinetics revealed by EdU lineage tracing during regeneration

**DOI:** 10.1101/627851

**Authors:** Bradley Pawlikowski, Nicole Dalla Betta, Tiffany Antwine, Bradley B. Olwin

**Author notes:** To whom correspondence should be addressed and current address: Bradley Olwin Ph.D., University of Colorado, Boulder, Department of Molecular, Cellular and Developmental Biology, 347 UCB, Boulder, CO 80309. Email addresses.

## Abstract

Skeletal muscle maintenance and repair is dependent on the resident adult muscle stem cell (MuSC). During injury, and in diseased muscle, stem cells are engaged to replace or repair damaged muscle, which requires the stem cells to exit quiescence and expand, followed by differentiation to regenerate myofibers and self-renewal to replenish the stem cell population. Following an injury, little is known regarding the timing of MuSC (skeletal muscle stem cell) self-renewal, myoblast expansion or myoblast differentiation. To determine the timing and kinetics of these cell fate decisions, we employed DNA-based lineage tracing to label newly replicated cells and followed cell fates during skeletal muscle regeneration. MuSCs activate and expand as myoblasts rapidly following injury, where the majority differentiate into myonuclei, establishing the centrally located myonuclear pool. Re-establishing the majority MuSC pool by self-renewal occurs after 5 days post-muscle injury, accompanied by low levels of myonuclear accretion that generate only peripheral myonuclei. In aged mice, possessing ∼1/2 the number of MuSCs present in young adult mice, the timing of post injury MuSC self-renewal is delayed, and although MuSCs expansion as myoblasts in aged muscle is impaired, the number of MuSC unexpectedly recovers to young adult levels during regeneration. Following an induced muscle injury, we found that myonuclei are generated within the first four days post injury derived from myoblasts expanding from activated MuSCs. Only later during regeneration, from 5 d to 14 d post injury, is the MuSC pool replenished by self-renewal, accompanied by generation of peripheral myonuclei.

## Introduction

Skeletal muscle’s ability to regenerate following injury is dependent on a resident muscle stem cell (MuSC) population (Lepper, Partridge, & Fan, 2011; Murphy, Lawson, Mathew, Hutcheson, & Kardon, 2011; Sambasivan et al., 2011). Individual, quiescent MuSCs in uninjured tissue occasionally divide and differentiate to maintain myofiber function (Pawlikowski, Pulliam, Betta, Kardon, & Olwin, 2015)(Keefe et al., 2015). In contrast, following an induced injury, MuSCs exit quiescence, undergo cell growth with an initial division between 24 and 48 hours post injury, and then divide rapidly, expanding as myoblasts (Webster, Manor, Lippincott-Schwartz, & Fan, 2016). Myoblast numbers peak around 4 days post injury and then gradually decline reaching steady state by 28 days post injury (Murphy et al., 2011)(Hardy et al., 2016). Despite a solid understanding of the molecular mechanisms that underlie differentiation, quiescence, and self-renewal, the timing of these events *in vivo* and the role of the environment are not known.

Muscle regeneration is diminished in aged mice, where repaired myofibers are smaller, generate less force, and muscle tissue is more fibrotic (Brack et al., 2007; I M Conboy et al., 2005; Cosgrove et al., 2014; Hall, Banks, Chamberlain, & Olwin, 2010; Mann et al., 2011; Zhang et al., 2016). MuSC function in aged mice is impaired, with a delayed first division, slower initial expansion, impaired self-renewal and enhanced differentiation (Bernet et al., 2014; Irina M Conboy, Conboy, Smythe, & Rando, 2003; Cosgrove et al., 2014; Elabd et al., 2014; Li, Han, Cousin, & Conboy, 2015; Shefer, Van de Mark, Richardson, & Yablonka-Reuveni, 2006). The decline in MuSC function with age contributes to impaired muscle regeneration in aged mice and is driven by cell autonomous and non-cell autonomous mechanisms (Bernet et al., 2014; Chakkalakal, Jones, Basson, & Brack, 2012; I M Conboy et al., 2005; Cosgrove et al., 2014; Dumont et al., 2015; He et al., 2013; Kuswanto et al., 2016; Liu et al., 2018; Price et al., 2014; Sacco et al., 2010; Tierney et al., 2014).

To assess the timing for myonuclear production and MuSC self-renewal *in vivo*, we used DNA base-labeling and lineage tracing to determine that all MuSCs enter the cell cycle after injury, that all centrally localized myonuclei are generated within 4 d post injury, and that the majority of MuSC self-renewal occurs after day 5 post injury, when additional myonuclear accretion generates peripherally-localized myonuclei. In aged mice, possessing ∼ 50% of adult MuSCs, post injury MuSC self-renewal is delayed and despite slower MuSCs expansion, the number of MuSCs unexpectedly is restored to adult numbers late during regeneration.

## Results

We initially asked whether or not all MuSC enter the cell cycle following an induced muscle injury, and whether the resultant myoblasts proliferate to produce myonuclei. To label MuSCs that enter the cell cycle, mice were provided EdU in the drinking water starting one day prior to muscle injury. The Extensor digitorum longus muscle (EDL) muscle was injured by BaCl_2_ injection and mice were maintained on EdU water until collection at 14 days post injury. MuSCs on isolated EDL myofibers at 14 d post injury, were identified by Pax7 immunoreactivity and by ClickIT chemistry to visualize EdU incorporation. On regenerated myofibers, identified by the presence of centrally located nuclei, 100% of the MuSCs and myonuclei were EdU+ (Fig 1A). Thus, following an induced injury all MuSCs enter the cell cycle, consistent with our prior published data (Troy et al., 2012), and differentiation of myonuclei is preceded by cell division (Fig 1A).

**Figure 1.**
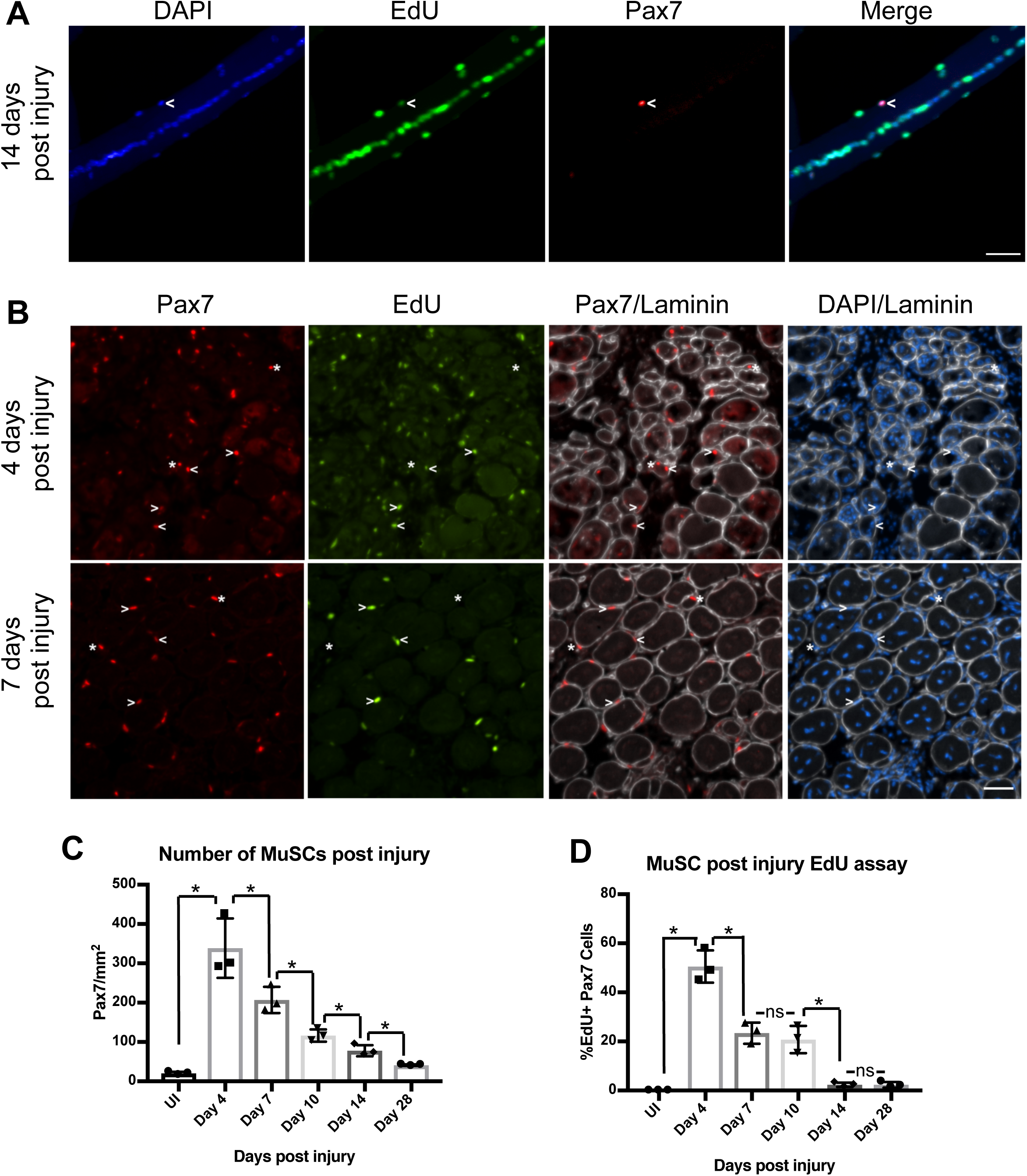
All MuSCs divide following injury and cease dividing by day 14 post injury. (A) Images of 14 d post injury EDL myofibers processed to detect EdU incorporation (green) and for Pax7 immunoreactivity to label MuSCs (red). DAPI is blue. White carets identify examples of EdU+/Pax7+ MuSCs in all figures (B) Post injury TA (tibialis anterior) muscle sections treated to visualize EdU+ cells (green), immunoreactivity for laminin (white), or immunoreactivity for Pax7 (red). Asterisks in all figures identify examples of EdU-/Pax7+ MuSCs. (C) Quantification of MuSCs at the indicated days post injury (n=3). (D) Quantification of EdU incorporation by MuSCs at indicated days post injury (n=3). Statistical significance was tested using Student’s t test. Asterisks indicates P-value < 0.05. NS indicates a P-value > 0.05. Scale bar is 40 μm.

To assess MuSC division kinetics following a muscle injury, we collected BaCL_2_ injured tibilias anterior (TA) muscles at various days post injury (4 d, 7 d,10 d, 14 d, and 28 d), giving 3 injections of EdU, 2 hours apart, immediately prior to collection. MuSCs in tissue sections from the injured TA muscles were identified by Pax7 and laminin immunoreactivity and then processed for EdU incorporation (Fig 1B; Supplemental Information Fig S1). Once activated following injury, Pax7+ cells divide rapidly, increasing over 20-fold by 4 d post injury then gradually decline to within 2-fold of the initial MuSC number by 28 d post injury (Fig 1C). Without injury, we detected no Pax7+/EdU+ cells following the 6h EdU exposure prior to muscle harvest (Fig 1D). The highest percentage of Pax7+/EdU+ cells was detected at 4 d post injury (∼50%), which declined to 20% at 7 d, and remained at 20% at 10 d post injury (Fig. 1D). The lowest percentage of Pax7+/Edu+ cells was detected at 14 days post injury with no further decrease by day 28 (Fig 1D). We conclude that self-renewal of MuSCs occurs within 14 d post injury and that all replenished MuSCs arise from a prior cell division.

To refine the timing of post injury differentiation and self-renewal we employed a variety of pulse-chase times to label nuclei with EdU. Mice were given EdU in the drinking water for the entirety of the regeneration (0-14 d), for the first 4 d post injury (0-4 d) or for days 5-14 post injury (5-14 d) followed by collection of EDL muscles at 14 days post injury (Schematic in Fig 2A). EdU incorporation and Pax7 immunoreactivity were quantified from individual myofibers with centrally located nuclei, using Pax7 immunoreactivity to identify MuSCs, and counting myonuclei as Pax7 negative cells located inside the myofiber. All myonuclei and all MuSCs are EdU+ when EdU is provided in the drinking water for the entirety of the injury/regeneration (0-14 d) (Fig 2B, E). In contrast, when EdU is administered from 0 d to 4 d post injury, a minority of MuSCs (∼20%) are EdU+, while all centrally located myonuclei, and some peripheral myonuclei are EdU+ (Fig 2C, E). Once EdU is incorporated into the DNA by cell division, additional cell divisions in the absence of EdU will dilute the EdU, which will eventually become undetectable. Therefore, the EdU+ MuSCs and the EdU+ centrally located myonuclei divided only a few times prior to self-renewal or differentiation, respectively. We conclude that most early MuSC cell divisions are devoted to myonuclei production as nearly all differentiated myonuclei are labeled within the first four days post-muscle injury.

**Figure 2.**
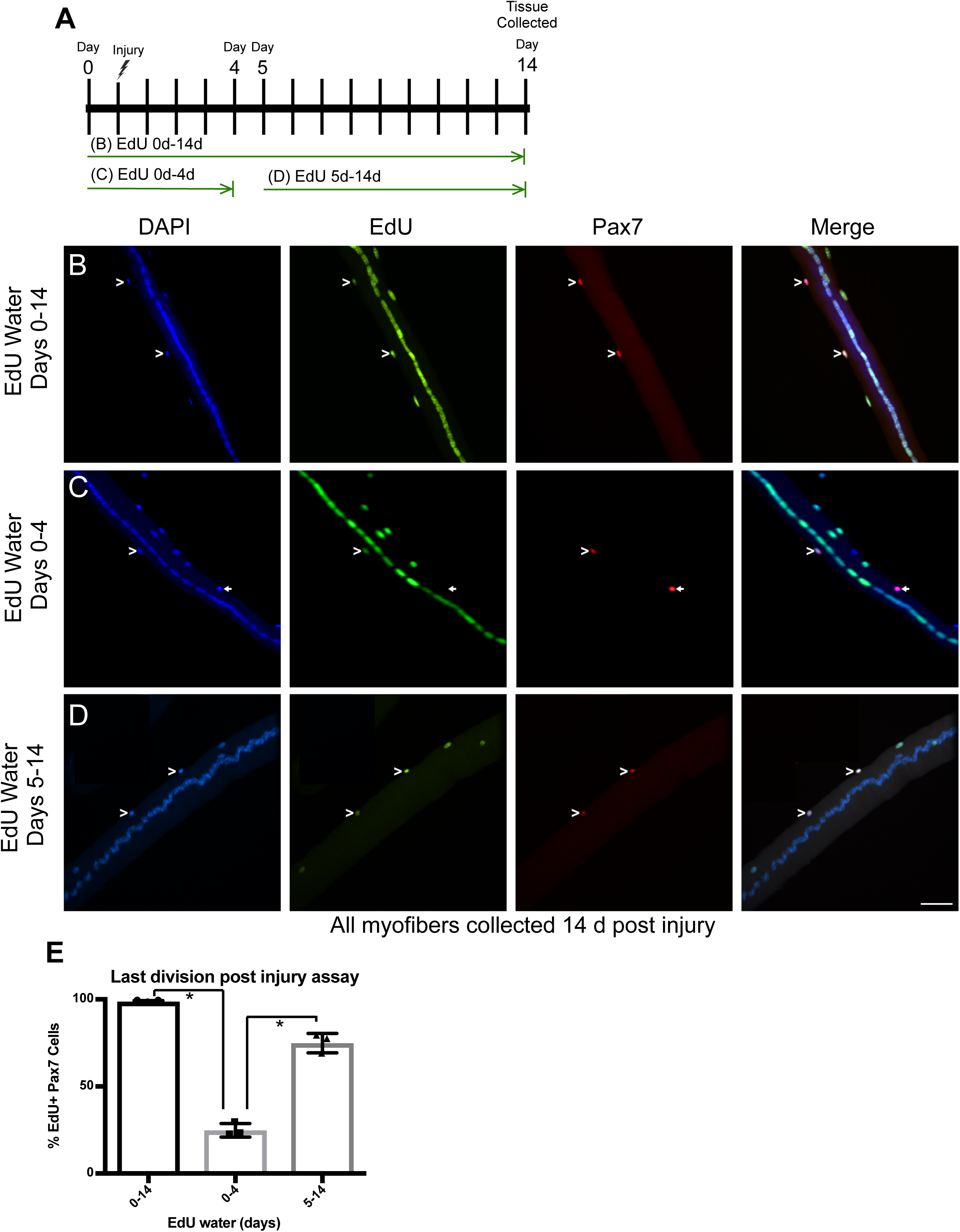
Kinetics of post injury MuSC division and differentiation. (A) Schematic of experimental design. (B-D) Images of 14 d post injury EDL myofibers, collected after the indicated exposures to EdU, and processed to detect EdU incorporation (green) and for Pax7 immunoreactivity to label MuSCs (red). DAPI is blue. White carets mark EdU+ MuSCs. White arrows mark EdU negative MuSCs. (B) EdU water given from day 0-14 post injury. (C) EdU water given from day 0-4 post injury. (D) EdU water given from day 5-14 post injury. (E) Quantification of EdU+ MuSCs on myofibers with central nuclei (n=3). Statistical significance was tested using Student’s t test. Asterisks indicates P-value < 0.05. Scale bar is 40 μm.

The majority of MuSCs are EdU+ when EdU is administered from 5 d to 14 d post injury and while some peripheral myonuclei are EdU+, no centrally localized myonuclei are EdU+, (Fig 2D and 2E). The majority of MuSCs undergo their final cell division between 5 d and 14 d post injury and self-renew but do not contribute to centrally localized myonuclei. Instead, from 5 d to 14 d post injury the products of self-renewal are MuSCs and peripherally localized myonuclei.

To further refine the time courses for myonuclear production and MuSC self-renewal following an induced muscle injury, we administered EdU for shorter durations. We provided EdU water from 0-2 d, 5-7 d, 8-10 d and 10-12 d post muscle injury, harvesting myofibers from the EDL muscles at 14 d post injury (Schematic in Fig 3A). On post injury myofibers, exposed to EdU water from day 0-2, less than 1% of MuSCs are EdU+ and centrally located myonuclei are weakly labeled (Fig 3B and 3E). Since MuSCs go through their first division between 24 and 48 hours (Jones et al., 2005; Troy et al., 2012; Webster et al., 2016), few if any MuSCs self-renew occurs during the first cell division following injury (Fig 3E). Furthermore, the weak EdU signal seen in the central nuclei is consistent with a MuSC dividing a few times after Edu removal at 2 d post injury as myoblasts prior to terminally differentiating (Fig 3B). Approximately 75% of MuSC self-renewal occurs between 5 d and 12 d post injury with 44% self-renewing between 5 d and 7 d post injury, 22% self-renewing from 8 d to 10 d post injury, and 6% self-renewing between 10 d and 12 d post injury, respectively (Fig 3C-E). MuSC self-renewal is accompanied by the appearance of peripherally located EdU+ myonuclei suggesting the peripheral nuclei arise via asymmetric cell division during MuSC self-renewal (Fig3 B-D).

**Figure 3.**
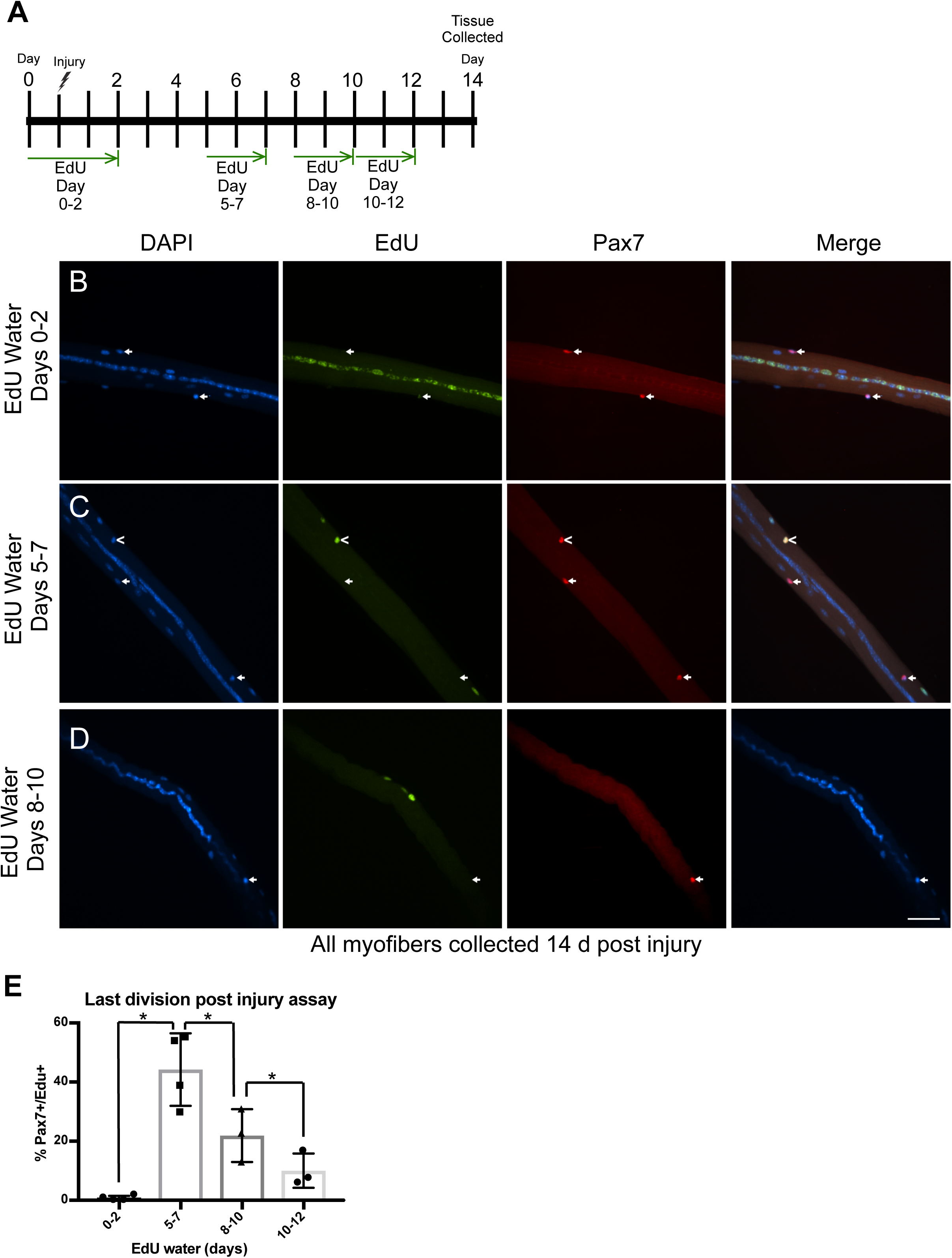
Kinetics for MuSC self-renewal. (A) Schematic of experimental approach. (B-D) Images of 14 d post injury EDL myofibers, collected after the indicated exposures to EdU water, and processed for EdU incorporation (green) and Pax7 immunoreactivity (red). DAPI is blue. White carets mark EdU+ MuSCs. White arrows mark EdU– MuSCs. (B) EdU water was given from day 0-2 post injury. (C) EdU water was given from day 5-7 post injury. (D) EdU water was given from day 8-10 post injury. (E) Quantification of EdU+ MuSCs on myofibers with central nuclei (n ≥ 3). Student’s t test was used for statistical analysis. Asterisks indicates P-value < 0.05. Scale bar is 40 μm.

### MuSC behavior following a muscle injury in aged muscle

MuSC function in aged muscle is impaired, (Bernet et al., 2014; Cosgrove et al., 2014; Price et al., 2014; Tierney et al., 2014) but whether aging affects the timing of post injury MuSC expansion, differentiation, or self-renewal is not known. We performed EdU labeling assays in aged (24-28 mo old) mice as detailed for adult mice. We collected BaCl_2_ injured TA muscles following 3 injections of EdU given at 2 h intervals and analyzed muscle sections at various time points post injury, for Pax7 immunoreactivity and EdU incorporation. In agreement with prior reports, MuSC numbers were significantly reduced in the un-injured aged TA muscle compared to adult TA muscle, young 20 ± 2 MuSCs/mm^2^ vs aged 8 ± 2 MuSCs/mm^2^ (Fig 4D and Supplemental Information Fig S2) (Brack, Bildsoe, & Hughes, 2005; Chakkalakal et al., 2012; Collins, Zammit, Ruiz, Morgan, & Partridge, 2007; Gibson & Schultz, 1983; Shefer et al., 2006; Sousa-Victor et al., 2014; Zhang et al., 2016). MuSCs from aged muscle expanded more slowly than in adult muscle (Fig 4D), but in contrast to adult muscle, MuSCs in aged muscle increase from 4 to 7 days post injury as opposed to decreasing (Fig 4A-4D). The resultant numbers of Pax7+ cells in aged and adult muscle are similar from 7 to 28 d post injury (Fig 4D), despite fewer Pax7+/EdU+ cells in aged muscle at all time points compared to adult muscle, respectively (Fig 4E). Although the number of MuSCs are reduced pre-injury in aged mice, the numbers of MuSCs in aged and adult muscle at 28 d post injury are indistinguishable (Fig 4D, 4E; Supplemental Information Fig S2).

**Figure 4.**
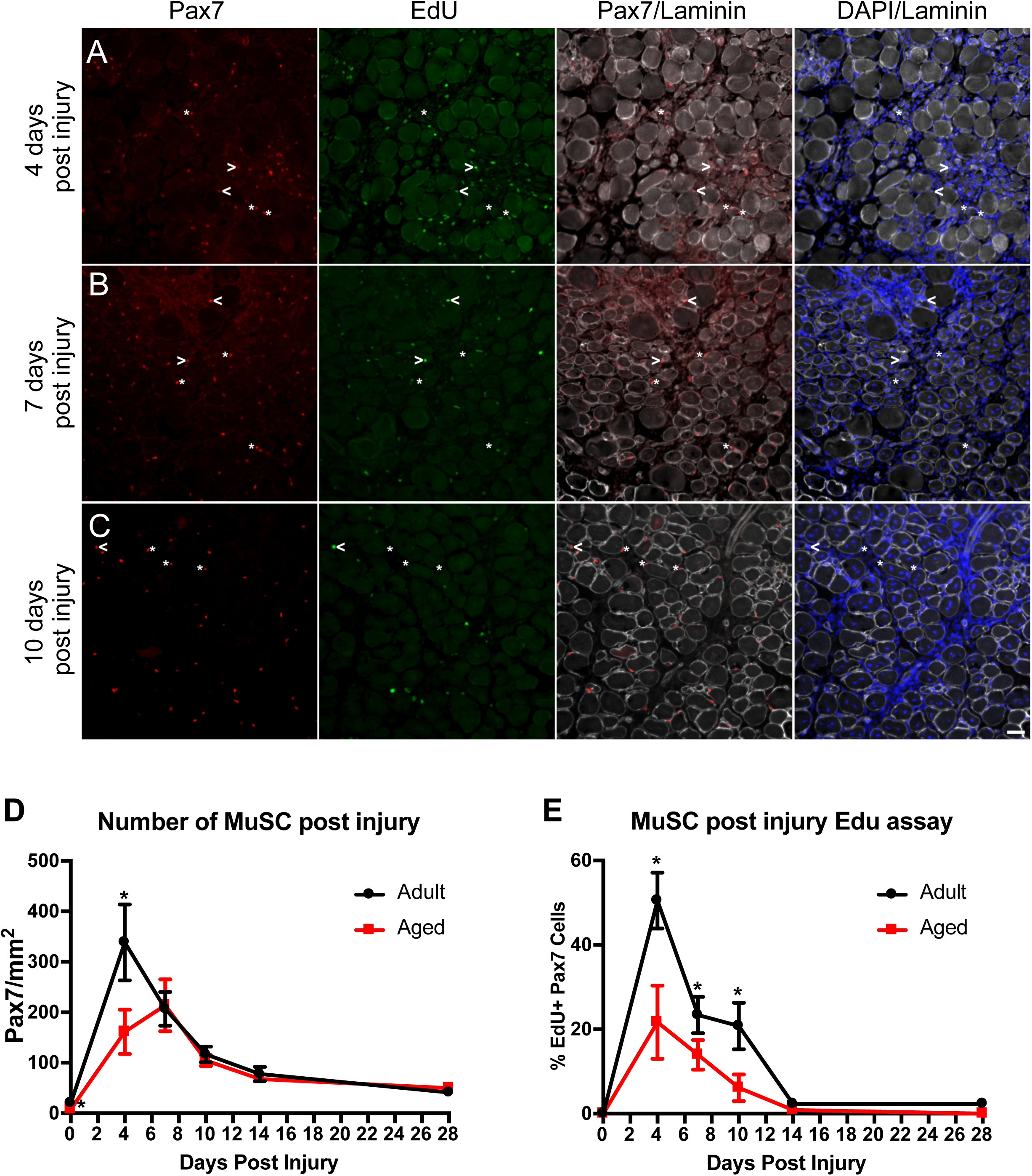
Kinetics for MuSC expansion in aged muscle. (A-C) Post injury TA muscle sections were processed for EdU detection (green), for laminin immunoreactivity (white), and for Pax7 immunoreactivity (red). White carets mark examples of EdU+ MuSCs. White asterisk mark examples of EdU negative MuSCs. (D) Quantification comparing the number of MuSCs in young adult versus aged tissue at various days post injury (n=3). (D) Quantification of EdU incorporation in MuSCs comparing young adult with aged muscle tissue at the indicated days post injury (n=3). Statistical significance was tested using Student’s t test. Asterisks indicates P-value < 0.05. NS indicates a P-value > 0.05. Scale bar is 40 μm.

Since the expansion of myoblasts in aged mice is altered compared to adult mice, we asked if the timing for MuSC self-renewal was similarly changed. Aged mice were administered EdU in their drinking water and the EDL muscles injured. The EdU water was administered either for the first 4 d post injury or from 5 d to 14 d post injury (Schematic in Fig 5A). We scored EdU incorporation and Pax7 immunoreactivity on the 14 d post injury myofibers with centrally located nuclei (Fig. 5B, 5C). When EdU water was given from 0 d to 4 d post injury only 10-15% of MuSCs in aged muscle self-renew, a ∼50% reduction compared to adult muscle (Fig5 5B, 5D). In contrast the number of MuSCs that self-renew between 5 d and 14 d post injury were similar in aged muscle compared to young muscle (Fig 5C, 5E). The timing of MuSC self-renewal is altered in aged muscle compared to adult muscle yet in both aged and adult mice MuSC self-renewal appears complete by 14 d post injury.

**Figure 5.**
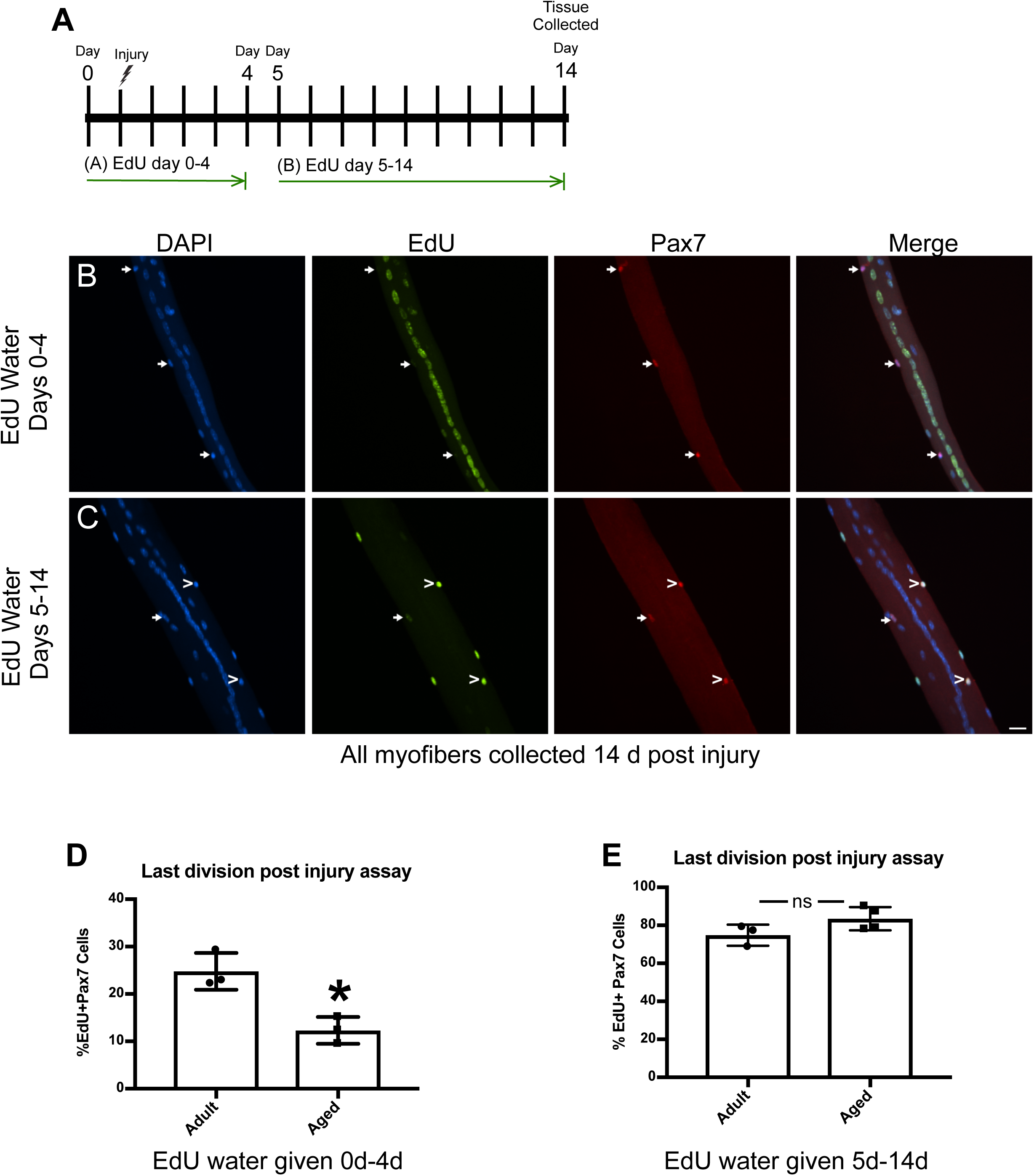
Kinetics for MuSC self-renewal in aged muscle. (A) Schematic of experimental approach. (B-C) Images of 14 d post injury EDL myofibers, collected after the indicated exposures to EdU water, and processed for EdU detection (green) and or Pax7 immunoreactivity (red). DAPI is blue. White carets mark EdU+ MuSCs. White arrows mark EdU negative MuSCs. (B) EdU water given from day 0-4 post injury. (C) EdU water was given days 5-14 post injury. (D) Quantification comparing the percent EdU+ MuSCs on 14 d post injury myofibers with central nuclei that were given EdU water from day 0-4 post injury in adult versus aged mice (n=3). (E) Quantification comparing the percent EdU+ MuSCs on 14 d post injury myofibers with central nuclei that were given EdU water from days 5-14 post injury in adult versus aged mice (n=3). Statistical significance was tested using Student’s t test. Asterisks indicates P-value < 0.05. Scale bar is 40 μm.

## Discussion

MuSC self-renewal has been extensively studies in culture, often using the first cell division on intact myofiber explants as a model (Kuang, Kuroda, Le Grand, & Rudnicki, 2007; Le Grand, Jones, Seale, Scimè, & Rudnicki, 2009; Troy et al., 2012). To compare the *in vitro* data to *in vivo* behavior, we asked when MuSC self-renewal and differentiation occurs post muscle injury in adult mice. We then asked if the kinetics for MuSC self-renewal and differentiation *in vivo* are distinct in adult vs. aged muscle. In contrast to intact myofiber cultures where MuSC self-renewal is studied during the first cell division, the majority of MuSC self-renewal occurs between day 5 and day 7 post injury *in vivo* and is complete by 14 d post injury. In aged mice, the reduced numbers of MuSCs prior to injury fail to expand as efficiently as MuSCs from adult mice and exhibit altered timing for self-renewal. However, in both young adult and aged mice, MuSC differentiation occurring prior to day 4 post injury produces centrally located myonuclei, while the majority of MuSC self-renewal and peripheral myonuclei are generated after 5 d post injury.

### Post injury muscle stem cell behavior in adult mice

Following an induced muscle injury with either BaCl_2_ or snake venom, skeletal muscle morphology does not return to pre injury status until 28 d to 60 d post injury (Hardy et al., 2016). Despite our finding that all MuSCs enter the cell cycle following an induced muscle injury, MuSC division and self-renewal are complete by day 14 post injury. Since the MuSCs present at day 14 post injury are the self-renewed population, we identified that the majority of post injury MuSC self-renewal occurs from 5-14 d post injury with the majority of MuSCs self-renewing between days 5-7 post injury, and the remaining completed by 10 d post injury. The last post injury cell division is likely the division that self-renews the satellite cell pool where an asymmetric division generates one quiescent MuSC daughter and one myonucleus. Consistent with this possibility, MuSC division rates are constant from day 7 to day 10 post injury despite a decrease in overall MuSC numbers. Future experiments to determine the precise timing for MuSC G0 entry and quantification of asymmetric cell divisions *in vivo* will be difficult but will be required to further support our hypothesis.

A prediction of self-renewal via asymmetric division is the generation of one MuSC daughter renewing the population and one daughter myoblast that can either proliferate or terminally differentiate. A striking feature of the EdU pulse assays was the exclusive generation of peripheral myonuclei at 5 d or later post injury, while prior to 5 d post injury the vast majority of myonuclei are centrally located. We propose that the location of the MuSC at the time of the last division results in the distinct myonuclear localization. During the first few days of muscle regeneration, MuSC division occurs within regenerating fibers (Webster et al., 2016), and the initial wave of MuSC proliferation gives rise to almost entirely all centrally located myonuclei. After day 5 post injury MuSCs are located on outside the plasma membrane of the myofiber (Fig 1B) and generate peripheral myonuclei that we predict are a product of asymmetric cell divisions to repopulate the MuSC niche.

An adult EDL myofiber contains approximately 250-300 myonuclei and has on average 5-6 MuSCs (Amthor et al., 2009; Bruusgaard, Liestøl, Ekmark, Kollstad, & Gundersen, 2003; Pawlikowski et al., 2015; White, Biérinx, Gnocchi, & Zammit, 2010). All myonuclei in a regenerated myofiber are produced from the initial 5-6 MuSCs present prior to injury. We demonstrate that the majority of myonuclei are generated by 4 d post injury and given the first MuSC division occurs 36 h post injury, and subsequent divisions occur every 12 h (Troy et al., 2012; Webster et al., 2016), then approximately 6 MuSC population doublings can occur from 36hr to 96hr (4 days) post injury. Therefore, the 5-6 MuSCs/EDL myofiber can yield approximately 300 differentiated myonuclei by 96 hr post injury, but only if self-renewal (and return to quiescence) was very limited. Our data fit this model (see Graphical Abstract Fig 6) where we propose that the majority of MuSCs must expand to generate the requisite myonuclei to rebuild muscle tissue, followed later by MuSC self-renewal to replenish the stem cell pool. Of particular interest is to determine whether the ∼20% of MuSCs that appear to self-renew and/or avoid differentiation prior to 4d post injury are a result of stochastic events or are a distinct population driven by a cell intrinsic program. Additionally, determining whether a fraction or the majority or peripherally generated myonuclei arise from asymmetric divisions and involved in MuSC self-renewal will aid in understanding the mechanisms responsible.

**Figure 6.**
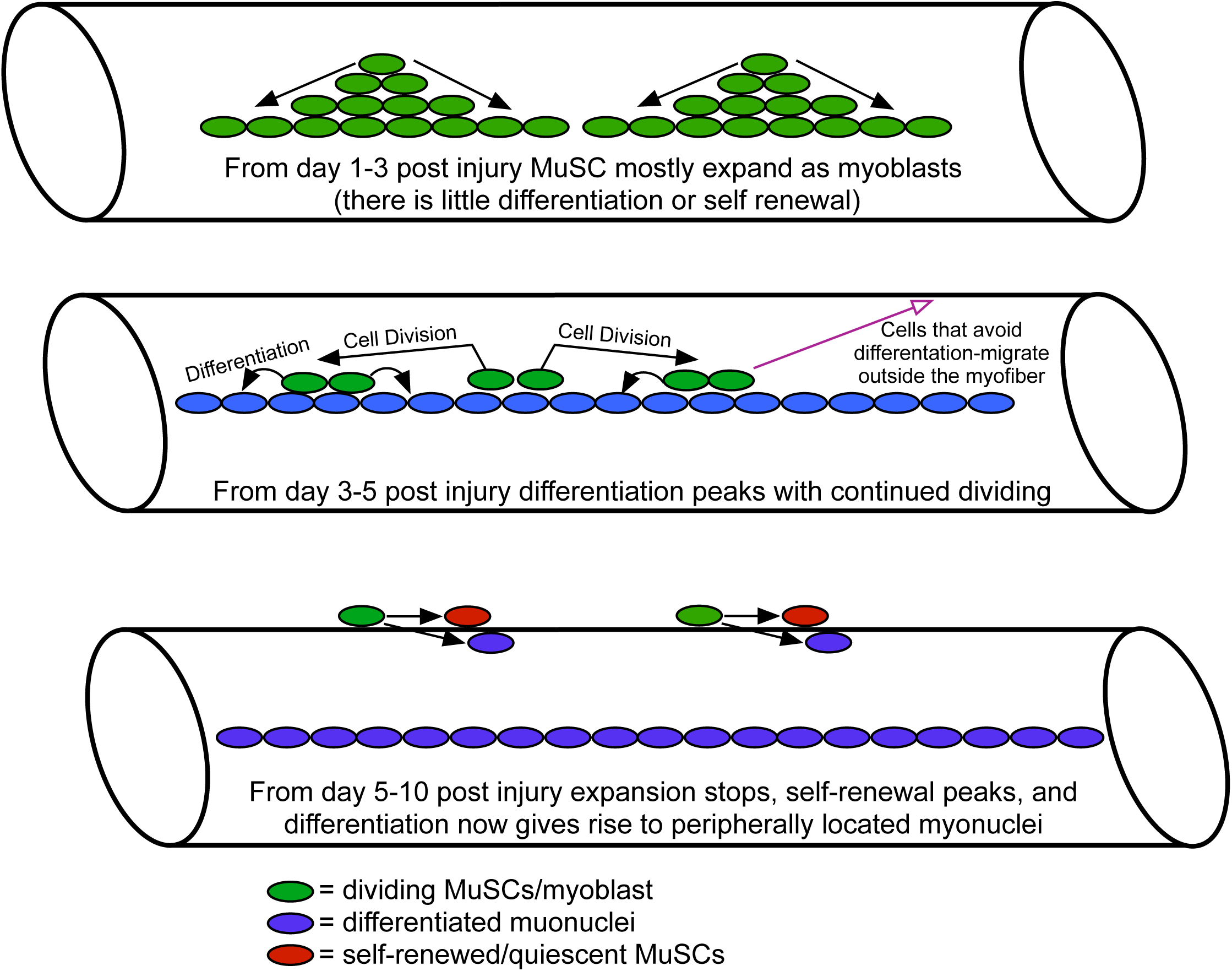
Graphical abstract depicting the timing of post injury MuSC expansion, differentiation and self-renewal.

### Post injury muscle stem cell behavior in aged mice

Skeletal muscle in aged mammals loses function, is reduced in mass, possess smaller myofibers and reduced numbers of MuSCs (Brack et al., 2005; Chakkalakal et al., 2012; Collins et al., 2007; Gibson & Schultz, 1983; Shefer et al., 2006; Sousa-Victor et al., 2014; Zhang et al., 2016). Part of the impairment of skeletal muscle with age is due to cell intrinsic deficits (Bernet et al., 2014; Cosgrove et al., 2014; Price et al., 2014; Tierney et al., 2014) and thus, the failure of skeletal muscle in aged mammals to regenerate likely arises from MuSC deficits (Bernet et al., 2014; Chakkalakal et al., 2012; Cosgrove et al., 2014; Liu et al., 2018; Pisconti et al., 2016; Zhang et al., 2016). To test if aging affects post injury MuSCs self-renewal or myonuclei generation we used EdU lineage tracing to compare MuSCs behavior in adult and aged mice.

The number of MuSC numbers in un-injured aged muscle is significantly reduced compared to adults (Fig. 4) (Brack et al., 2005; Chakkalakal et al., 2012; Collins et al., 2007; Gibson & Schultz, 1983; Shefer et al., 2006; Sousa-Victor et al., 2014; Zhang et al., 2016) and MuSC expansion to generate centrally located myonuclei following a muscle injury in aged tissue is impaired compared to adult mice. In adult mice, the maximum number of Pax7+ cells are achieved by 4 d post injury and then declines, while in contrast, MuSCs continue to expand from 4-7 d post injury in regenerating aged muscle. Is the expansion of MuSCs in aged muscle from day 4 to day 7 post injury occurring at the expense of myonuclear generation and MuSC self-renewal? At 4 d post injury there is a 2-fold reduction in Pax7+/Edu+ cells, which could contribute to smaller myofiber size and inefficiencies in regeneration. Unexpectedly, at 14 d post injury in aged muscle, MuSC numbers exceed those originally present in uninjured aged tissue, restoring the numbers that were present as an adult. Because self-renewal is impaired in aged MuSCs as compared to adult MuSCs, it is likely that the newly generated MuSCs in aged muscle will be incapable of an additional round of regeneration following a second injury. MuSC expansion and self-renewal are altered *in vivo* in aged muscle tissue compared to young tissue following a muscle injury, suggesting the altered kinetics contribute to impaired muscle regeneration as muscle ages.

We have developed an EdU lineage assay to robustly assess the kinetics of myonuclear production and MuSC self-renewal following a muscle injury in adult and aged skeletal muscle. Unexpectedly, we found that immediately following an injury the majority of cell divisions are devoted to producing myonuclei, while the majority of MuSCs self-renew later, between 5d and 7 d of regeneration. In injured, aged muscle, aberrant MuSC expansion as compared to young adult muscle fails to provide sufficient numbers of myonuclei that may lead to reductions in myofiber size as well as inefficient regeneration. The disruption of the kinetics or a reduction in MuSC cell function as observed in aging will further impair regeneration as all MuSCs are engaged to repair muscle tissue following a large muscle injury and may be in part responsible for the increased fibrosis and fat deposition observed in aged skeletal muscle.

### Experimental Procedures

#### Mice

Mice were bred and housed according to National Institutes of Health (NIH) guidelines for the ethical treatment of animals in a pathogen-free facility at the University of Colorado at Boulder. University of Colorado Institutional Animal Care and Use Committee (IACUC) approved animal protocols and procedures. Young mice used in figures 1-3 were C57B6 mice, (Jackson Labs Stock No. 000664) between 4 and 8 months old and were a mix of male and female. The aged mice used in figures 4 and 5 mice were F1 mice from a C57BL/6J and DBA/2J cross (Jackson Labs No. 100006), collected between 24-28 months old and a mix of male and female.

#### Mouse Injuries and Edu delivery

Mice were anesthetized with isofluorane followed by injected with 50μL of 1.2% BaCl_2_ into the left TA and EDL muscle. To deliver EdU, mice were either given water containing 0.5mg/ml EdU (Carbosynth), with 1% glucose or given IP injections of 10 mM EdU (Carbosynth), re-suspended in water, a volume of 100 µL per 25g mouse weight.

#### Myofiber isolation, immunostaining and culture

The EDL muscles were dissected and placed into 400 U/mL collagenase at 37°C for 1.5h with shaking and then transferred into Ham’s F-12C and 15% horse serum to inactivate the collagenase. Individual EDL myofibers were separated using a glass pipet and immediately fixed using 4% paraformaldehyde for 10 min at room temperature and stored in PBS at 4°C. To detect Pax7 and for visualization of EdU incorporation, myofibers were permeabilized with 0.5% Triton-X100 in PBS (phosphate buffered saline without calcium or magnesium pH 7.4), containing 3% bovine serum albumin (Sigma) for 30 min at room temperature. EdU incorporation was visualized using the Click-iT EdU Alexa Fluor 488 imaging kit (ThermoFisher) following the manufacturer’s protocol. The anti-Pax7 antibody, used at 2 μg/mL (Developmental Studies Hybridoma Bank (DSHB) at the University of Iowa) was incubated with intact myofibers at room temperature for 1h followed by three washes in PBS and then myofibers were incubated with and donkey anti-mouse Alexa Flour 555 (ThermoFisher) secondary antibody for 1hr at room temperature. Myofibers were washed 3 times in PBS, incubated with 1 μg/mL DAPI for 10 min at room temperature to label nuclei and then mounted in Mowiol supplemented with DABCO (Sigma-Aldrich) as an anti-fade agent.

#### Immunofluorescence staining of tissue section

The TA muscle was dissected, fixed for 2h on ice cold 4% paraformaldehyde, and then transferred to PBS with 30% sucrose at 4°C overnight. Muscle was mounted in O.C.T. (Tissue-Tek®) and cryo-sectioning was performed on a Leica cryostat to generate 8 μm sections. Tissues and sections were stored at −80°C until staining. Tissue sections were post-fixed in 4% paraformaldehyde for 10 minutes at room temperature and washed three times for 5 min in PBS. To detect Pax7 on muscle sections we employed heat-induced epitope retrieval. Post-fixed sections on slides were placed in a citrate buffer (100mM Sodium citrate containing 0.05% Tween20 at pH 6.0) and subjected to 6 min of high pressure-cooking in a Cuisinart model CPC-600 pressure cooker. For immunostaining, tissue sections were permeabilized with 0.5% Triton-X100 (Sigma) in PBS containing 3% bovine serum albumin (Sigma) for 30 min at room temperature. EdU was visualized as described for myofibers. Primary antibodies included anti-Pax7 (DSHB) at 2 μg/mL and rabbit anti-laminin (Sigma-Aldrich cat#L9393) at 2.5 μg/mL incubated on muscle sections for 1h at room temperature. Alexa secondary antibodies, donkey anti mouse Alexa Flour 647, donkey anti-rabbit Alexa Flour 555 (ThermoFisher) were used at a 1:750 dilution and incubated with muscle sections for 1 h at room temperature. Prior to mounting, muscle sections were incubated with 1 μg/mL DAPI for 10 min at room temperature then mounted in Mowiol supplemented with DABCO (Sigma-Aldrich) as an anti-fade agent.

#### Microscopy and image analysis

All images were captured on a Nikon inverted spinning disk confocal microscope. Objectives used on the Nikon were: 10x/0.45NA Plan Apo and 20x/0.75NA Plan Apo Images were processed using Fiji ImageJ. Confocal stacks were projected as maximum intensity images for each channel and merged into a single image. Brightness and contrast were adjusted for the entire image as necessary.

#### Statistics

Statistical analysis was performed in Prism (GraphPad), where statistical significance was assessed using, two-tailed, unpaired Student’s *t* test with *p* < 0.05 considered significant. Each N was generated from an individual mouse.

## Acknowledgements

This work was funded by grants from the ALSAM Foundation (BBO), the Muscular Dystrophy Association (BBO), the Glenn Foundation for Medical Research (BBO) and NIAMS AR070630 (BBO).

## Author Contributions

BP and BBO conceived the experiments. BP, ND, and TA performed the experiments analyzed the data, and made figures. BP and BBO wrote the manuscript. BBO supervised the research. All authors read and approved the manuscript.

Supplement Information Figure S1-Images of post injury tissue from adult mice

(A) TA muscle sections processed for EdU incorporation (green), for laminin immunoreactivity (white), and for Pax7 immunoreactivity (red). White carets mark examples of EdU+ MuSCs. White asterisk mark examples of EdU negative MuSC. Time points of tissue collection are indicated on left. Scale bar is 40 μm.

Supplement Information Figure S2-Images of post injury tissue from aged mice

(A) TA muscle sections stained for EdU incorporation (green), with an antibody to laminin (white), and with an antibody to Pax7 (red). White asterisk mark examples of EdU negative MuSC. Time points of tissue collection are indicated on left. Scale bar is 40 μm.

