## Supplemental Figures 1 and 2 for "Skeletal muscle stem cell self-renewal and differentiation kinetics revealed by EdU lineage tracing during regeneration"

Supplemental Information Figure S1- Images of post injury tissue from adult mice

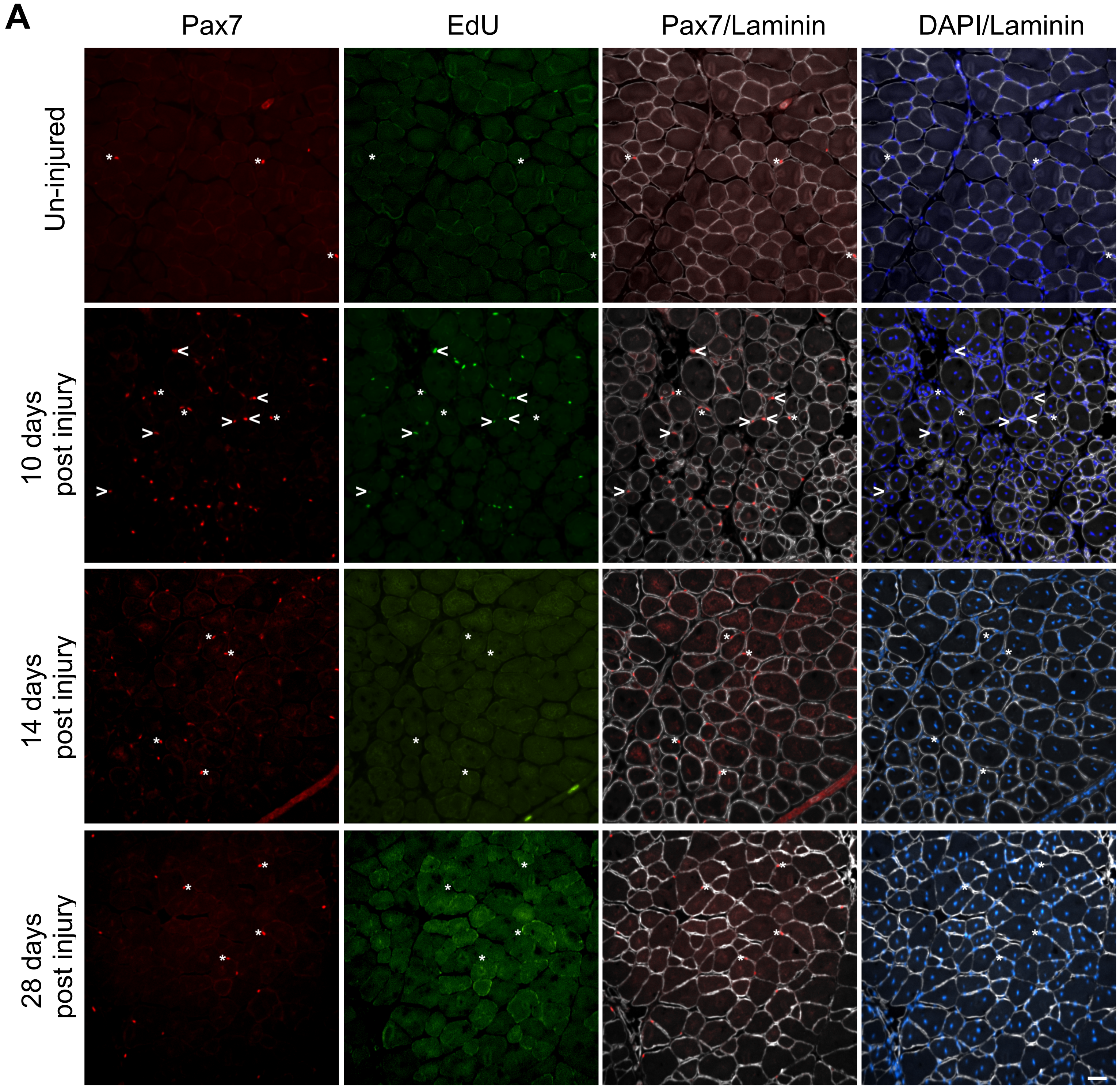

Suplemental Information-Figure S2 Images of post injury tissue from aged mice

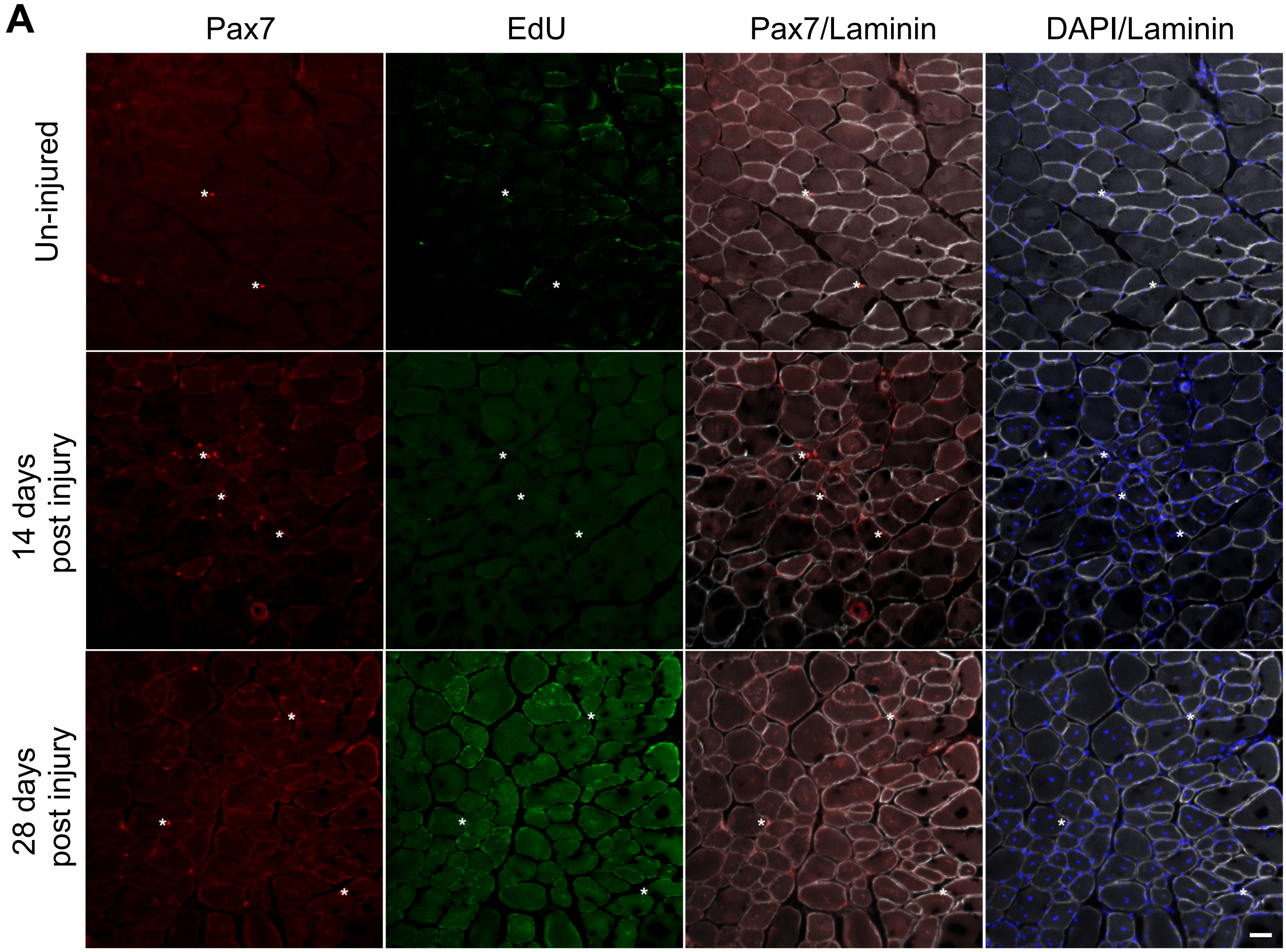
